# Epigenetic clock and methylation studies in elephants

**DOI:** 10.1101/2020.09.22.308882

**Authors:** Natalia A. Prado, Janine L. Brown, Joseph A. Zoller, Amin Haghani, Mingjia Yao, Lora R. Bagryanova, Michael G. Campana, Jesús E. Maldonado, Ken Raj, Dennis Schmitt, Todd R. Robeck, Steve Horvath

**Affiliations:** Center for Species Survival, Smithsonian Conservation Biology Institute, Front Royal, VA, USA; Center for Conservation Genomics, Smithsonian Conservation Biology Institute, Washington, D.C., USA; Department of Biostatistics, Fielding School of Public Health, University of California, Los Angeles, Los Angeles, California, USA; Department of Human Genetics, David Geffen School of Medicine, University of California, Los Angeles, Los Angeles, California, USA; Department of Epidemiology, Fielding School of Public Health, University of California, Los Angeles, Los Angeles, California, USA; Radiation Effects Department, Centre for Radiation, Chemical and Environmental Hazards, Public Health England, Chilton, Didcot, UK; College of Agriculture Missouri State University, SeaWorld Parks and Entertainment, Orlando, FL USA

**Author notes:** Joint Senior Author (Janine L. Brown, Todd R. Robeck, Steve Horvath). <u>Corresponding author</u> Steve Horvath, PhD, ScD Address: Gonda Building, 695 Charles Young Drive South, Los Angeles, CA 90095. <u>Emails:</u>.

**Keywords:** elephant, aging, development, epigenetic clock, DNA methylation

## Abstract

Age-associated DNA-methylation profiles have been used successfully to develop highly accurate biomarkers of age (“epigenetic clocks”) in humans, mice, dogs, and other species. Here we present epigenetic clocks for African and Asian elephants. These clocks were developed using novel DNA methylation profiles of 140 elephant blood samples of known age, at loci that are highly conserved between mammalian species, using a custom Infinium array (HorvathMammalMethylChip40). We present epigenetic clocks for Asian elephants (*Elephas maximus*), African elephants (*Loxodonta africana*), and both elephant species combined. Two additional human-elephant clocks were constructed by combing human and elephant samples. Epigenome-wide association studies identified elephant age-related CpGs and their proximal genes. The products of these genes play important roles in cellular differentiation, organismal development, metabolism, and circadian rhythms. Intracellular events observed to change with age included the methylation of bivalent chromatin domains, targets of polycomb repressive complexes, and TFAP2C binding sites. These readily available epigenetic clocks can be used for elephant conservation efforts where accurate estimates of age are needed to predict demographic trends.

## INTRODUCTION

In comparison to many other mammals, elephants have remarkably long lives, with African (*Loxodonta africana*) and Asian (*Elephas maximus*) elephant lifespans exceeding 70 years and 80 years, respectively ^1,2^. Like humans, elephants have large brains relative to their body size, are self-aware, slow-growing, and operate in a highly structured social system ^3–5^. However, they have a lower risk of developing cancers compared to humans, and also do not have a comparable period of reproductive senescence ^6–8^. Global elephant populations are threatened by poaching and habitat destruction, and so are the focus of intensive conservation efforts. Such initiatives rely heavily on accurately predicting demographic trends and population viability; which are important mitigation strategies that require a reliable estimate of age.

Currently, the number of elephants in a population are visually estimated and reported as a proportion of mature individuals in an estimated area. This method however, is prone to gross overestimation in long-lived species that mature slowly, such as elephants, where visual determination of “maturity” is subjective and difficult ^9,10^. Furthermore, the use of tooth-based age criteria for African elephants developed by Laws (1966)^11^ is also problematic as it is based on discovered jaw bones of unknown ages, assumes proportionate distribution of tooth sequences relative to actual tooth progression rates, and assumes a maximum lifespan of 60 years ^1^. Furthermore, having been revised numerous times with data from wild elephants ^3,12^, museum specimens ^13^, and animals in zoos ^14^, these estimates are still based on animals of uncertain ages, and often, of small sample sizes ^1^. Other ageing techniques for elephants have been developed using biological data such as eye lens weight ^15,16^, hind leg weight ^17^, foot dimensions, tusk length and circumference, shoulder height ^10,15,18,19^, dung bolus circumference ^20^. But again, much of these morphological data were accrued from culled individuals, and not living animals of known ages. Thus, currently employed methods to age individual elephants are highly subjective, often inaccurate, and difficult to carry out in the field; hence, no single technique has been adopted universally for conservation purposes ^10,21,22^. As such, there is a pressing need to develop an objective and accurate estimator of elephant age.

DNA methylation is one of the best characterized epigenetic modifications that affects the activity of a DNA segment without changing its sequence. In mammals, it plays an important role in biological processes such as silencing of transposable elements, regulation of gene expression, genomic imprinting, X-chromosome inactivation, carcinogenesis, and aging ^23^. It was previously observed that the degree of cellular DNA methylation is influenced by age ^24,25^. The significance and specificity of these alterations remained a source of speculation until the development of an array-based technology that permitted quantification of methylation levels of specific CpG positions on the human genome. With this advancement came the opportunity and insight to combine age-related methylation changes of multiple DNA loci to develop a highly accurate age-estimator (epigenetic clocks) for all human tissues ^26,27^. For example, the human pan-tissue clock combines the weighted average of DNA methylation levels of 353 CpGs into an age estimate that is referred to as DNA methylation age (DNAm age) or epigenetic age ^28^. Importantly, the difference between DNAm age and chronological age (referred to as “epigenetic age acceleration”) is predictive of all-cause mortality in humans, even after adjusting for a variety of known risk factors ^29–33^. The notion that epigenetic age acceleration may be indicative of health status was confirmed by multiple reports that demonstrated its association with a multitude of pathologies and health conditions including, but not limited to, cognitive and physical decline ^34^, Alzheimer’s disease ^35^, Down syndrome ^36^, Werner syndrome ^37^, HIV infection ^33^, Huntington’s disease ^38^, obesity ^39^, and menopause ^40^. Today, epigenetic clocks are regarded as validated biomarkers of human age and are employed in human clinical trials to measure the effects and efficacy of anti-ageing interventions ^26,41,42^. The hope of extending the benefits of these clocks to other animals was initially encouraged by the direct applicability of the human pan-tissue clock on chimpanzee DNA methylation profiles. This compatibility, however, could not be extended to other animals because of evolutionary genome sequence divergence ^28^. Hence, it is necessary to develop *de novo* epigenetic clocks specific to animals of interest, as for example what has been accomplished for mice ^43–48^. The availability of an epigenetic clock specific for elephants would indeed be welcomed, as current methods to estimate age are fraught with problems.

In general, molecular biomarkers of aging for elephants could find two major areas of application: 1) comparative studies in the biology of aging and disease; and 2) *in-situ* and *ex-situ* conservation efforts. Here, we present several ready-to-use DNA methylation-based age estimators (epigenetic clocks) for Asian and African elephants, the characteristics of age-related CpGs that constitute the elephant epigenetic clocks, and how they compare with those of humans.

## Results

### Epigenetic clocks

We obtained methylation profiles of DNA from the blood of 140 elephants (57 African and 83 Asian), ranging in age from 1.20 to 73.6 years, using a custom Infinium array (HorvathMammalMethylChip40), which consists of probes that recognize approximately 38,000 specific CpGs, with neighboring DNA sequences that are conserved between different species of the mammalian class (Table 1). Random forest predictors of elephant species and sex led to perfect predictive accuracy, i.e. the out-of-bag estimates of error were zero.

**Table 1.** Description of blood methylation data from elephants. N=Total number of samples per species. Number of females. Age: mean, minimum and maximum.

| Species | Latin Name | N | No.<br>Female | Mean<br>Age | Min.<br>Age | Max.<br>Age |
| --- | --- | --- | --- | --- | --- | --- |
| Asian elephant | *Elephas maximus* | 83 | 67 | 37 | 2.36 | 73.6 |
| African elephant | *Loxodonta africana* | 57 | 43 | 24.9 | 1.20 | 48.5 |

To arrive at unbiased estimates of the epigenetic clocks, we performed cross-validation analyses and obtained estimates of the age correlation R (defined as Pearson correlation between the age estimate, DNAm age and chronological age), as well as the median absolute error.

From these we obtained epigenetic clocks for: 1) Asian elephants; 2) African elephants; and 3) both elephant species combined. These epigenetic clocks utilize 38, 49 and 52 specific CpGs respectively. There are 7 CpGs that are shared between the Asian and African elephant clocks. The dual elephant species clock shares 13 age-related CpGs with African elephant clock, 21 CpGs with Asian elephant clocks and 5 CpGs with both elephant clocks.

We also constructed dual-species epigenetic clocks that can be applied directly to humans and elephants. These clocks were similarly derived, except that the training data set was constituted by DNA methylation profiles of human tissues and elephant blood, all of which were derived using HorvathMammalMethylChip40. From these, we generated two human-elephant clocks, each of which can accurately estimate ages of humans and elephants. The difference between the two dual species clocks lies in how age is reported. One reports results in terms of chronological age in years, while the other reports relative age, which is the ratio of chronological age to maximum lifespan of the respective species, and assumes values between 0 and 1. For example, the relative age of a 61-year-old human is 61/122.5 = 0.5. This ratio allows alignment and biologically meaningful comparison between species with different lifespans.

The dual elephant epigenetic clock, which was trained on DNA methylation profiles from both African and Asian elephant blood, exhibited a high correlation of R=0.96 with median error of 3.3 years (Figure 1A). Equally significant correlations were obtained for the African elephant (R=0.97, Figure 1B) and Asian elephant (R=0.96, Figure 1C) epigenetic clocks. Likewise, the human-elephant clock for chronological age was just as accurate when both species were analyzed together (R=0.97, Figure 1D) or when the analysis was restricted to elephant blood DNA profiles only (R=0.96, Figure 1E). Finally, strongly significant correlations also were obtained with the human-elephant clock for *relative age* regardless of whether the analysis was applied to both species (R=0.97, Figure 1F) or only to elephants (R=0.96, Figure 1G).

**Figure 1:**
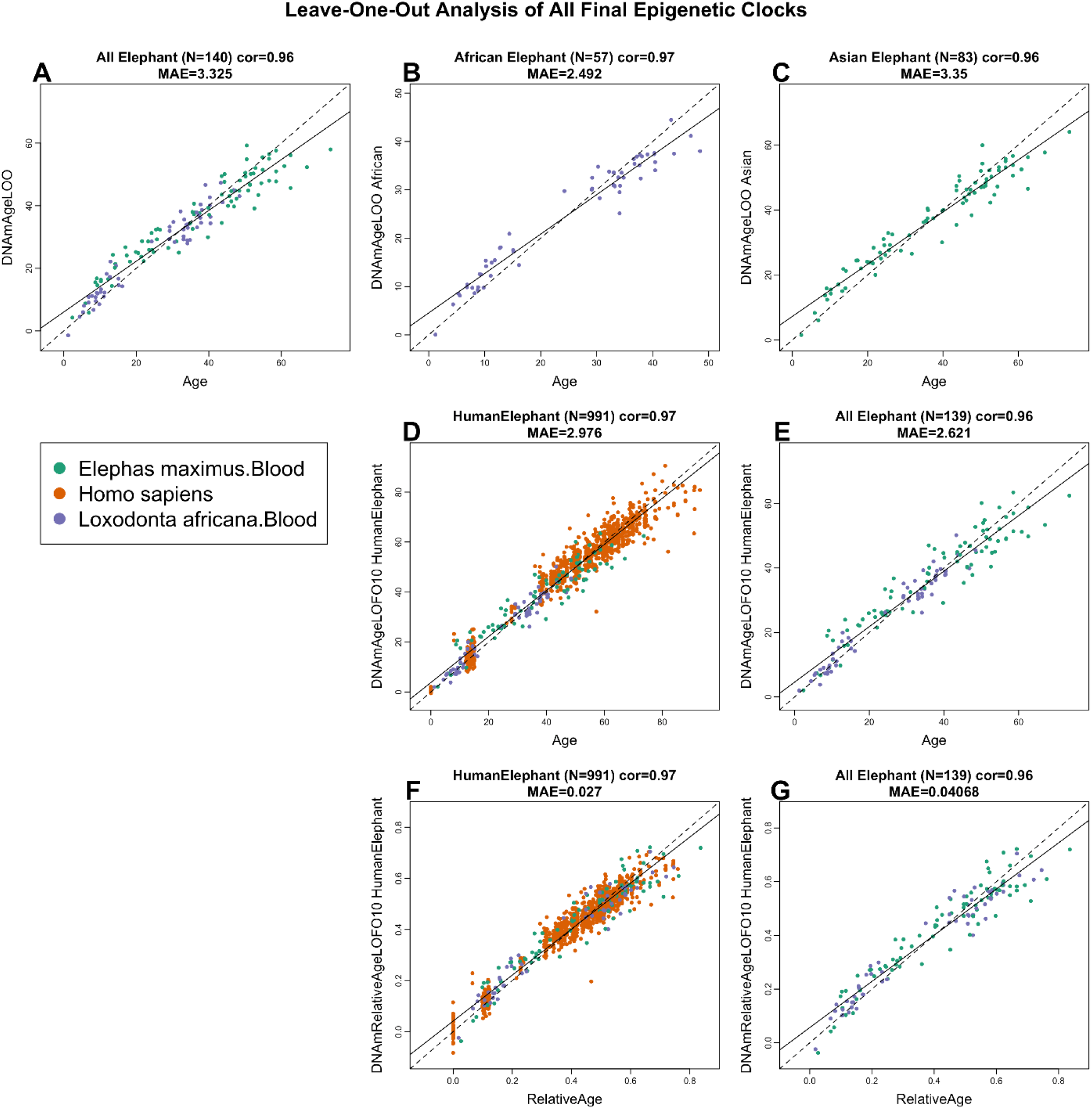
Cross-validation study of epigenetic clocks for African and Asian elephants and humans. Chronological age versus leave-one-sample-out (LOO) estimate of DNA methylation age (y-axis, in units of years) for epigenetic clocks for A) both African and Asian elephants combined, B) African elephant, C) Asian elephant. D) Ten fold cross validation analysis of the human-elephant clock for chronological age. Dots are colored by species (orange=human). E) Same as panel D) but restricted to elephants. F) Ten fold cross validation analysis of the human-elephant clock for relative age, which is the ratio of chronological age to the maximum lifespan of the respective species. G) Same as panel F) but restricted to elephants. Each panel reports the sample size, correlation coefficient, median absolute error (MAE).

### Characteristics of age-related CpGs

The successful derivation of the five epigenetic clocks that are applicable to elephants fulfils one of the aims of this endeavor, which was to identify biomarkers of elephant age and use them to create elephant age-estimating tools that are readily useable. The other aim was to uncover epigenetic features of the elephant genome that are associated with age that will allow for cross-species comparison with other members of the mammalian class. To do that, it was not sufficient to analyze only CpGs that constitute the elephant clocks, as they are just a subset of all age-related CpGs. Therefore, an epigenome-wide association study (EWAS) of elephant age was carried out using *Loxodonta africana*.loxAfr3.100 genome assembly, because a high quality genome assembly for the Asian elephant is presently unavailable. As the HorvathMammalMethylChip40 probes were selected based on highly conserved regions in mammalian genomes, the use of *Loxodonta africana*.loxAfr3.100 genome will not adversely affect the analysis and its applicability to the Asian elephant.

In total, 32,817 probes from the HorvathMammalMethylChip40 array could be aligned to specific loci that are proximal to 5,197 genes in the African elephant genome. Setting a genome-wide significance at p<10^−8^, the total number of CpGs whose methylation levels changed with age (age-related CpGs) were 775 for Asian and 1,298 for African elephants (Figure 2A). Epigenome wide-association studies (EWAS) identified 1,101 age-related CpGs that were shared between both the elephant species, and this translates to greater than 40% conservation (Figure 2C). Potential age-associated genes were identified by virtue of their proximity to the age-associated CpGs. The top age-related CpG of Asian elephants, is downstream of the PGM1 locus (correlation test Z statistic z = −14.3), while that of African elephants is upstream of the ZFHX3 gene (z = −14.5) (Figure 2A). Incidentally, this same CpG ranked second in the Asian elephant, making it and its proximal gene, ZFHX3 the highest-ranked between both the elephant species. Indeed, analysis of both elephants (meta-analysis) showed 8 CpGs upstream of ZFHX3 (Stouffer’s meta-analysis Z statistic = −8.7 to −17.9) to decrease methylation with age. Some other top hits in the meta-analysis included hypomethylation in ZNF536 intron, downstream region of FGF8, and downstream of CDK5RAP3. Although at the genome-wide level, a significant majority of age-associated CpGs are hypomethylated with age, at promoter regions they are primarily hypermethylated (Figure 2B). Analysis of CpGs in transcriptional factor binding-sites highlighted TFAP2C-binding sites in particular, as being hypermethylated with age, with some sites hypomethylated as well. Beyond identifying and considering potential age-associated genes and promoters individually, greater insights can be obtained by knowing the pathways, clinical outcomes and regulatory systems that are associated with these potential age-associated elements (genes and promoters). Such enrichment analysis identified cellular differentiation and development to be the major biological processes that are predicted to be affected by age-related CpGs (Figure 3). As for intracellular activities, the highest scores were seen with increased methylation at targets of polycomb repressive complex (SUZ12 and EED) and bivalent chromatin domain (H3K4me3 and H3K27me3). These features are shared between Asian and African elephants. Other important features that appear in only one of the elephant species are: genes involved in mTOR signaling, which become hypomethylated with age, and those that regulate Notch signaling and telomerase which become hypermethylated. These highlighted genes are notable for their reported involvement in ageing and will be discussed below.

**Figure 2.**
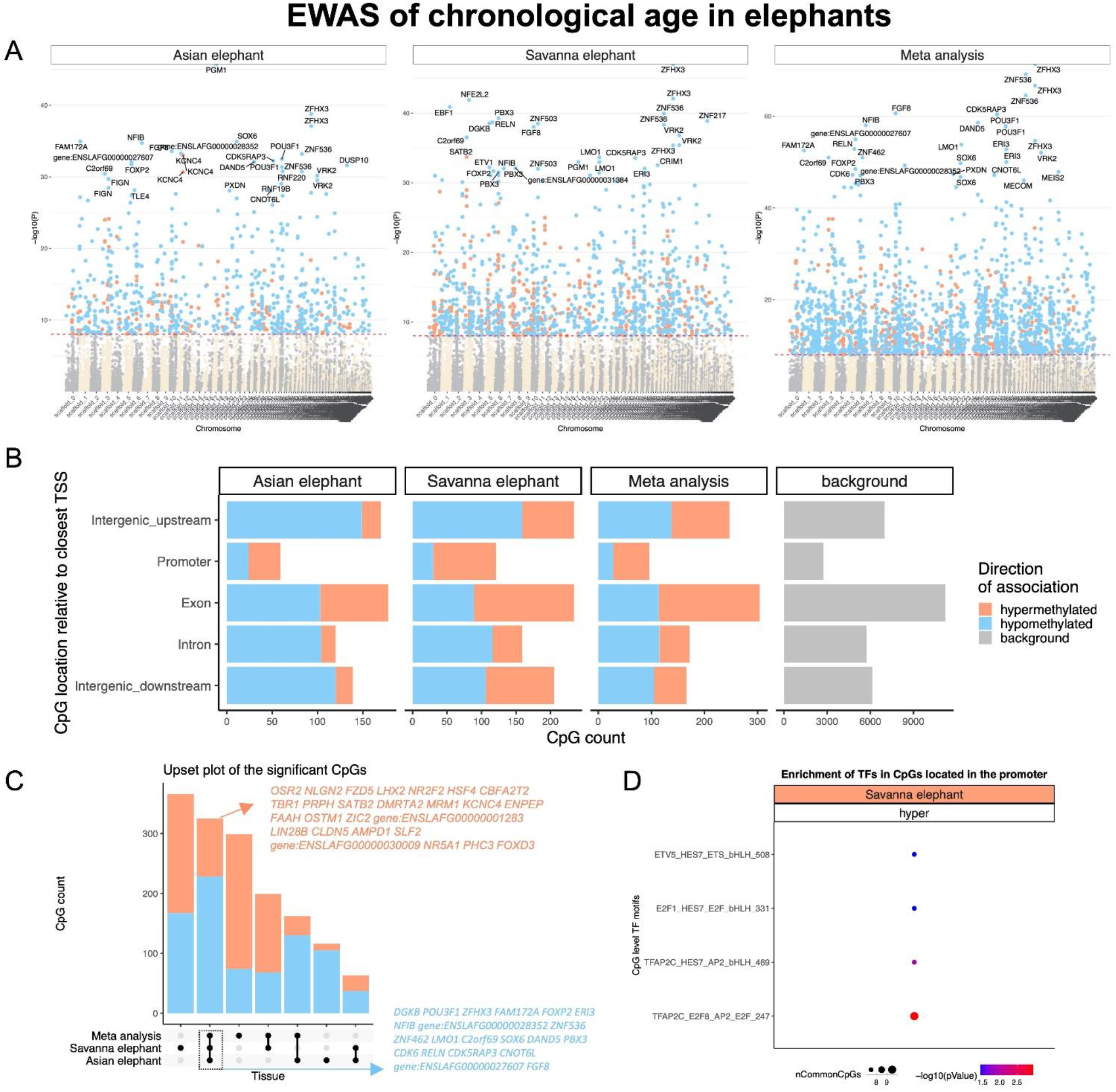
Epigenome-wide association (EWAS) of chronological age in blood of African elephants (Loxodonta africana) and Asian elephants (Elephas maximus). A) Manhattan plots of the EWAS of chronological age. The coordinates are estimated based on the alignment of Mammalian array probes to Loxodonta_africana.loxAfr3.100 genome assembly. The direction of associations with p < 10^−8^ (red dotted line) is highlighted by red (hypermethylated) and blue (hypomethylated) colors. Top 30 CpGs was labeled by the neighboring genes. B) Location of top CpGs in each species relative to the closest transcriptional start site. Top CpGs were selected at p < 10^−8^ and further filtering based on z score of association with chronological age for up to 500 in a positive or negative direction. The number of selected CpGs: Sevanna elephants, 953; Asian elephants, 666; and meta-analysis, 985. The grey color in the last panel represents the location of 32817 mammalian BeadChip array probes mapped to Loxodonta_africana.loxAfr3.100 genome. C) Upset plot representing the overlap of aging-associated CpGs based on meta-analysis or individual species. Around 20 Neighboring genes of the top overlapping CpGs were labeled in the figure. D) Transcriptional motif enrichment for the top CpGs in the promoter and 5’UTR of the neighboring genes.

**Figure 3.**
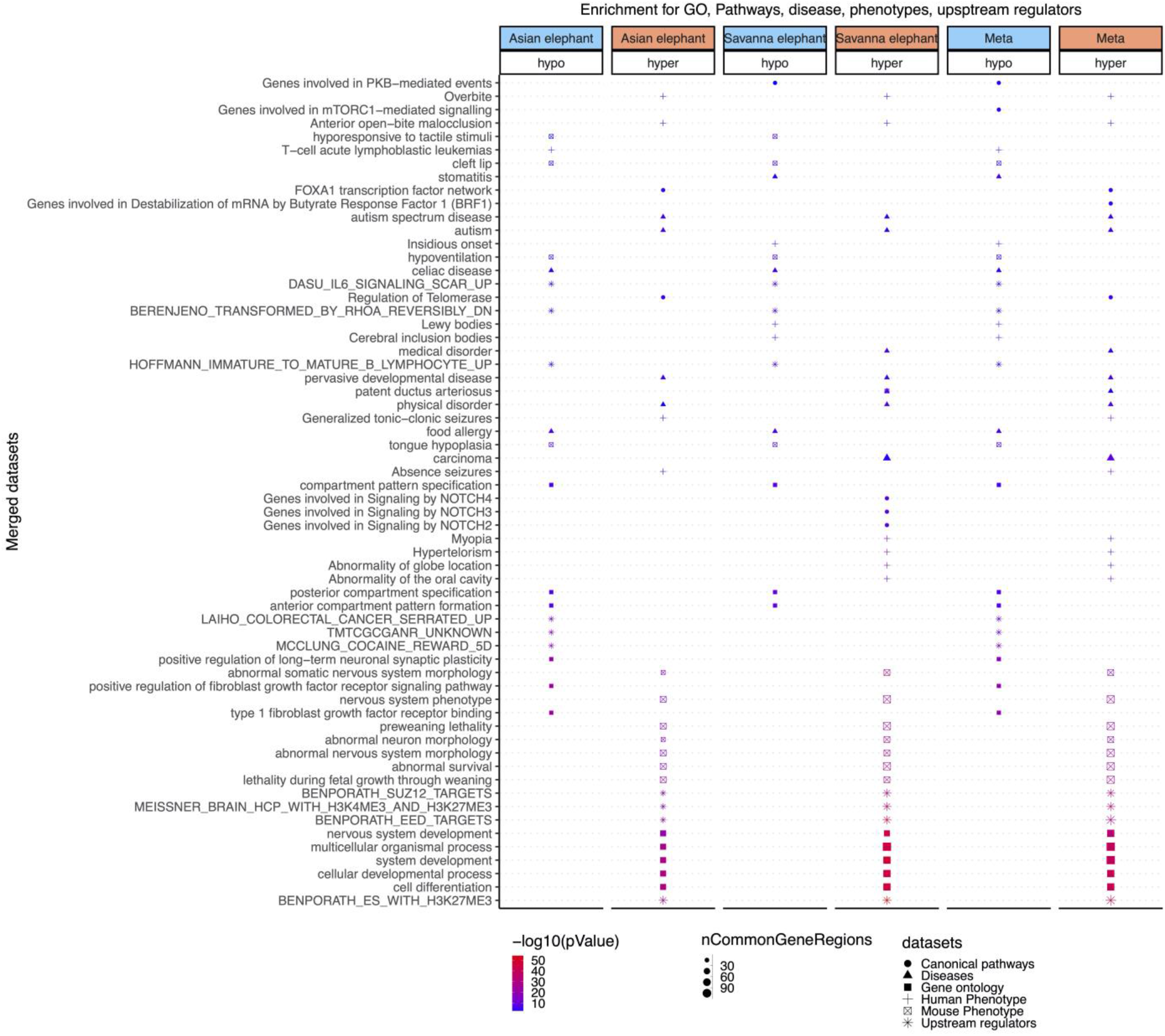
Enrichment analysis of the top CpGs associated with DNAm age in Asian, African elephants and meta-analysis. The analysis was done using the genomic region of enrichment annotation tool ^47^. The gene level enrichment was done using GREAT analysis ^47^ and human Hg19 background. The background probes were limited to 21,377 probes that were mapped to the same gene in the African elephant genome. The top three enriched datasets from each category (Canonical pathways, diseases, gene ontology, human and mouse phenotypes, and upstream regulators) were selected and further filtered for significance at p < 10^−4^.

### Comparison of DNAm aging between human and elephant

To allow a direct comparison between humans and elephants, which exhibit different lifespans, we converted the values of chronological age to relative age (age / maximum lifespan) for each species. Multivariate models were used to identify age-related CpGs that are either shared between humans and elephants, or distinct to either of the elephant species. At the 5% false discovery rate (FDR), the analysis identified 852 age-related CpGs (608 hypermethylation and 244 hypomethylation) that are share between humans and elephants (Figure 4A). In contrast, there were respectively, 64 and 83 age-related CpGs that are unique to the African and Asian elephant with respect to humans, and of these, more than 30% were shared between both elephant species. Some of the top age-associated CpGs shared between humans and elephants are those that are associated with ZFHX3, which was also the top identified gene from EWAS of elephants alone. The activity of this protein and its potential implication is discussed below.

**Figure 4.**
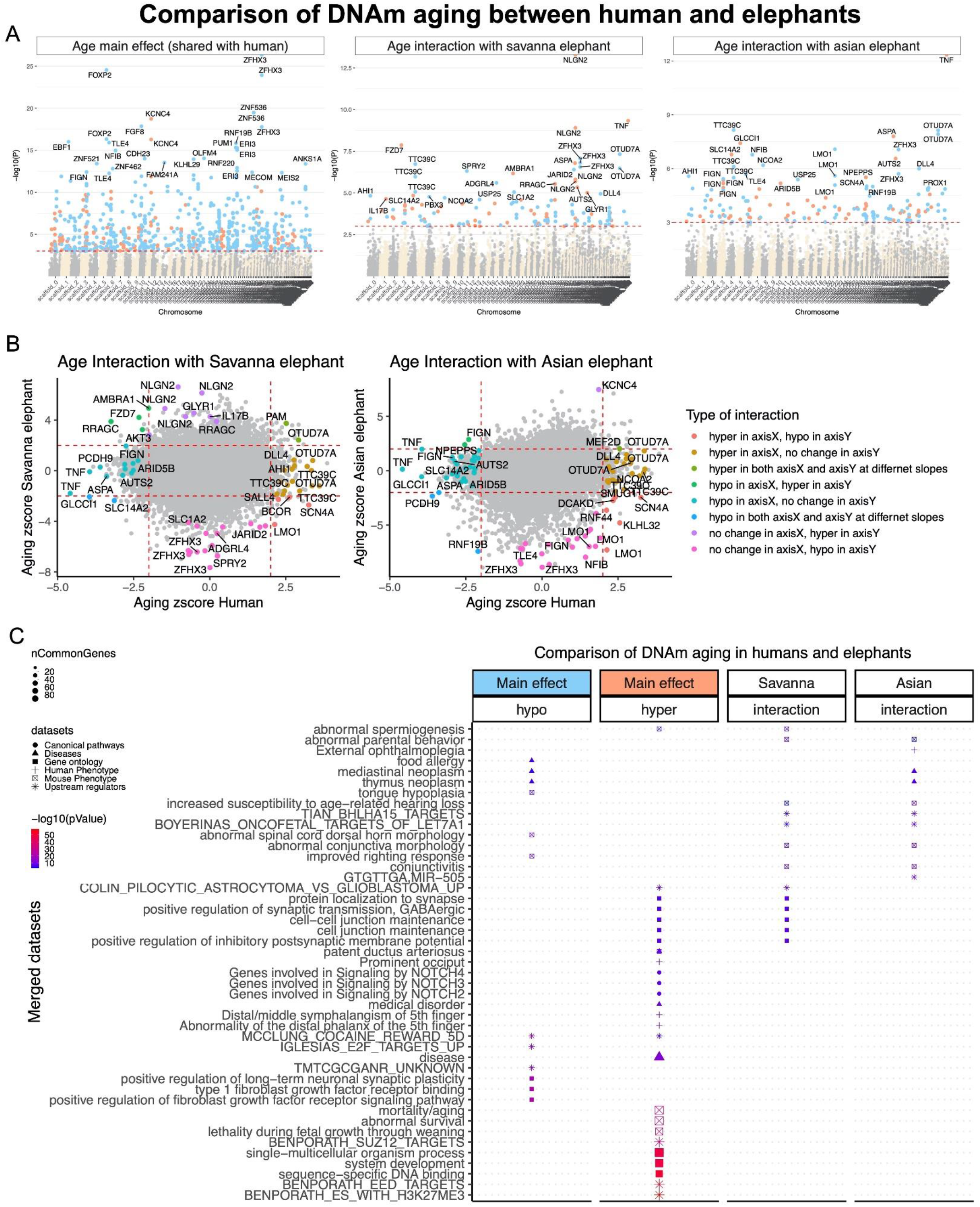
Comparison of DNAm aging between humans and elephants’ blood. A) Manhattan plots of the DNAm aging loci that are shared (main effect of aging) or interact between humans and elephants. The analysis is done by multivariate regression models with or without (to estimate the main effect) interaction term for relative age and species. The outcome in the models was DNAm levels. The analysis is limited to 21788 probes that are conserved between humans and elephants. Relative age is estimated by dividing chronological age per maximum reported lifespan (Humans: 122.5, African elephant: 65, Asian elephant: 88 years). Elephant samples were limited to the relative age of 0.2-0.65 to match the human samples. Sample size: Humans, 32; African elephant, 33; Asian elephant, 60. The coordinates are estimated based on the alignment of Mammalian array probes to Loxodonta_africana.loxAfr3.100 genome assembly. The red line in the Manhattan plot indicates p <1e-3. B) Scatter plots DNAm aging between humans and elephants. The highlighted CpGs are the loci with significant interaction between species at a 5% FDR rate. In total, nine categories of interaction were defined based on aging zscore of each species. C) Enrichment analysis of the genes proximate to CpGs with an aging association that are shared (aging main effect) or interact between humans and elephants. The gene-level enrichment was done using GREAT analysis ^74^ and human Hg19 background. The background probes were limited to 21,377 probes that were mapped to the same gene in the African elephant genome.

Some of the top age-related CpGs that are exclusive to elephants reside in the exon and promoter of the tumour necrosis factor (TNF) gene. As TNF-α is a major pro-inflammatory cytokine whose level increases with human age and is also associated with numerous human chronic diseases^49,50^, its exclusion from the human age-associated CpG was somewhat surprising. We nevertheless proceeded to overlay the elephant-specific age-related CpGs on scatterplots of methylation changes of age-associated CpGs between humans and elephants. This resulted in the identification of nine categories of age-related CpGs in elephants that are not shared with humans (**Figure 4B**). From this, the divergence with regards to TNF was more clearly observed, where CpGs proximal to TNF are hypomethylated with age in humans, but they either remained unchanged or are significantly hypermethylated in African and Asian elephants. Other examples of age-associated CpGs with stark divergent pattern between humans and elephants include: SCN4A (hyper in human, hypo in both elephant species), which is a sodium channel, and RRAGC (unchanged in human, hyper in African elephants), which promotes intracellular localization of the mTOR complex.

While the above identification of specific genes proximal to age-related CpGs can be informative, they can as well be confusing (as exemplified by TNF above) as biological effects are rarely, if ever elicited by the product and activity of a single gene. Instead, it is the cumulative effects of multiple genes within a pathway that generate a biological outcome. Therefore, it can be more informative to analyze the entire collection of implicated genes, for common pathways in which they are involved. We performed such enrichment analysis of genes proximal to age-associated CpGs that are either shared between elephants and humans, or distinct to either of the elephant species (**Figure 4C**). The age-related CpG proximal genes shared between humans and elephants enriched pathways related to development (e.g. NOTCH signaling), mortality, ageing and the nervous system. Furthermore, the analysis showed highly significant methylation of DNA targets of polycomb repressor complex (SUZ12 and EED targets) and H3K27me3 binding sites.

In contrast, elephant age-related CpGs that are not shared with humans are enriched with pathways that are involved in synapse, hearing, conjunctivitis, parental behavior, and also neoplasm (Figure 4C). Our analysis also underscored BHLHA15 and microRNA LET7A1 as potential age-associated upstream regulators of age-associated genes in elephants.

### Sex-associated DNA methylation of African and Asian elephants

We further investigated potential sex-associated differences in DNAm in both African and Asian elephants. The analysis focused on CpGs with baseline sex differences after adjusting for chronological age. At p< 10^−4^, African and Asian elephants had 898 and 972 differentially methylated CpGs between sexes (sex-related CpGs), respectively (**Figure 5A**). Interestingly, these sex-related CpGs were clustered in only a few chromosomes, and more than 70% were shared between African and Asian elephants (**Figure 5B**), with the top CpGs residing in the upstream region of the BCL6 Corepressor (BCOR). It is highly likely that some of these clusters are loci on the sex chromosomes. In general, sex-related CpGs were distributed in gene regions (**Figure 5C**). As expected, enrichment analysis of genes proximal to sex-related CpGs highlighted features associated with X-chromosome and X-linked inheritance (**Figure 5D**). Interestingly, some cognitive and behavioral differences such as mental health, spatial learning, coordination, and contextual conditioning emerged to be potentially associated with sex.

**Figure 5.**
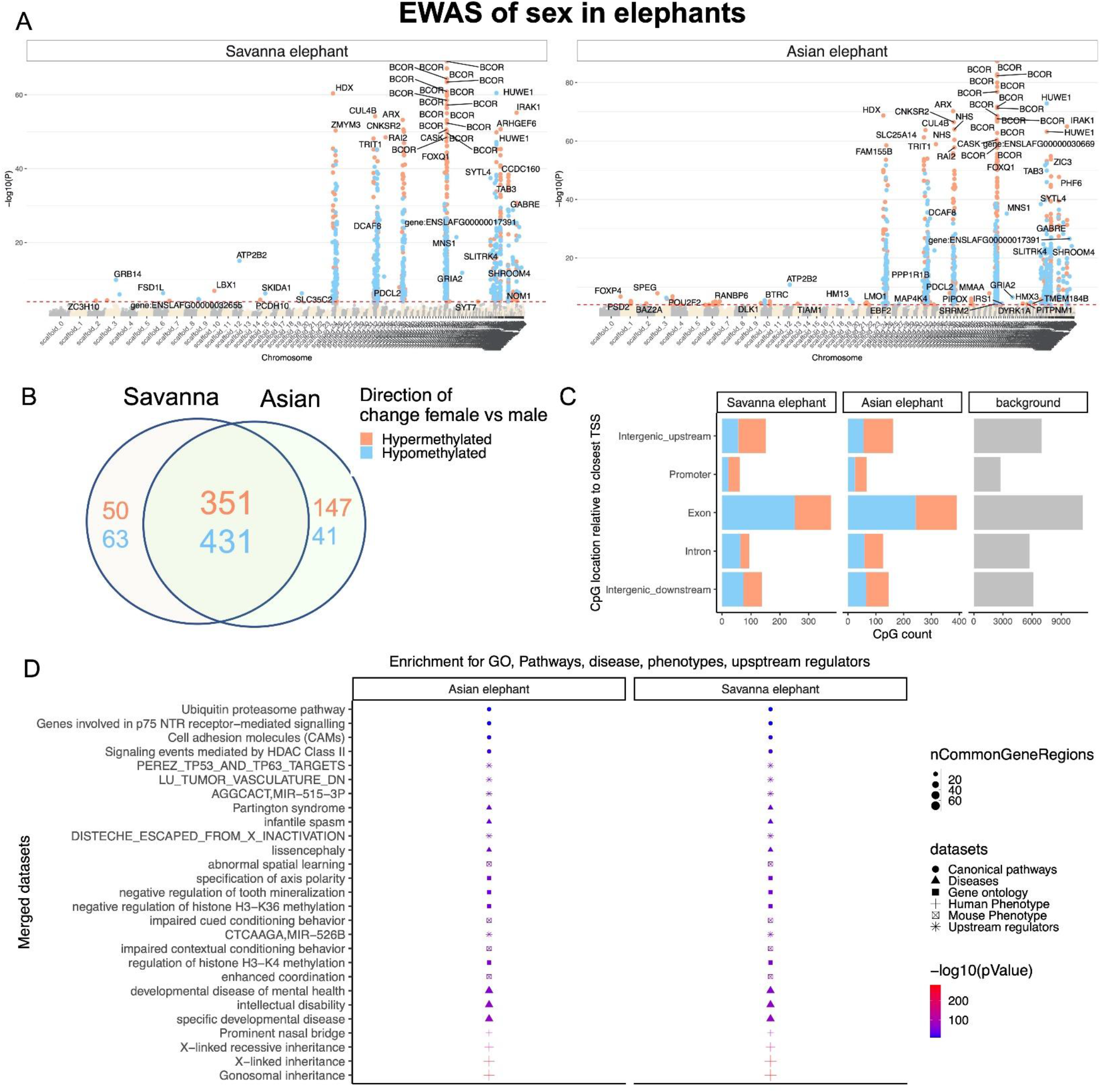
EWAS of sex difference in the blood of African and Asian elephants. A) Manhattan plots of sex differences in elephant samples. The association is estimated from a multivariate regression model with adjustment for chronological age in each species. The coordinates are estimated based on the alignment of Mammalian array probes to Loxodonta_africana.loxAfr3.100 genome assembly. The red line in the Manhattan plot indicates p <1e-4. B) Venn diagram of the sex-related differentially methylated CpGs in African and Asian elephants. The CpGs were selected at p<1e-4 significance level. C) Location of top CpGs in each species relative to the closest transcriptional start site. D) Enrichment analysis of the genes proximate to sex-related differentially methylated CpGs in elephants. The gene-level enrichment was done using GREAT analysis ^74^ and human Hg19 background. The background probes were limited to 21,377 probes that were mapped to the same gene in the African elephant genome.

## Discussion

This study describes five epigenetic clocks for elephants, of which three are pure elephant clocks (for Asian, African, and both species combined) and two are dual species human/elephant clocks that are applicable to humans as well. The human-elephant clocks for absolute and relative age demonstrate the feasibility of building epigenetic clocks for different species based on a single mathematical formula. This further consolidates emerging evidence that epigenetic aging mechanisms are conserved, at least between members of the mammalian class. Despite this, it was not a foregone conclusion that DNA methylation changes could be encapsulated by a single mathematical formula to accurately estimate age in the two disparate species (human and elephant). The critical step to removing the species barrier was the development and use of a bespoke DNA methylation array that profiles 38,000 CpGs with flanking DNA sequences that are conserved across numerous mammalian species. This allowed us to directly analyze DNA methylation profiles from different species, either individually or collectively, as they were all derived using the same DNA methylation measurement platform. With this unique platform, we profiled whole blood DNA samples from elephants of known ages in North American zoos, and were able to construct highly accurate epigenetic clocks that apply to the entire life course (from birth to old age) of both Asian and African species.

The most direct practical application of these readily useable epigenetic clocks will be in elephant conservation efforts. Accurate age determination and population statistics are of great importance to obtain an understanding of the health of herds, their maturity, sustainability and turn-over within a region. The current age estimation methods are prone to substantial errors as they are subjective and based on less-than-ideal criteria described in the introduction. With the immediate availability of these clocks, less than a milliliter of blood is required to obtained DNA methylation profile from the HorvathMammalMethylChip40 array. The resulting profiles need only be provided to the pre-set formula for the generation of elephant age. Apart from the standard techniques for blood collection and DNA extraction, no specialized skills, techniques or expensive equipment are required for their immediate deployment.

Another element of conservation is sex identification of elephant remains. While osteological and morphometric assessments can be used, they require good preservation state of the remains. Alternative DNA-based methods are sought, and recently, one that uses the ratio of X chromosome DNA sequence reads over autosomal DNA sequence read (Rx ratio method) was presented and shown to work even with ivory samples ^51^. Alternatively, one could use our DNA methylation based estimator of elephant sex.

There is increasing evidence that the process of ageing may be broadly similar between species of the mammalian class. This, however, does not preclude the possibility that in some species, specific pathways and modifications may operate simultaneously, and in parallel with the underlying basic mechanism of ageing. In this regard, it is notable that in humans and both elephant species, decreasing methylation of CpGs near the ZFHX3 locus were most significantly associated with age. The protein encoded by this gene is a transcription factor that is required to set the pace and amplitude of circadian rhythm ^52^. In humans, demethylation of this gene locus is augmented in those whose health are adversely affected by desynchronization of their circadian rhythm due to long-term night-shift work ^53^.

The GREAT enrichment analysis of the elephant EWAS of age implicated regulation of telomerase. Although not shared by the human data set analyzed here, the hTERT locus was previously identified as one of the strongest hits in EWAS of accelerated ageing of human blood ^54^. hTERT maintains telomere length, which determines proliferative capacity of cells. This in turn impacts cellular regeneration, which is linked to ageing. Despite this, the relationship of hTERT with epigenetic ageing is far from clear, as we have previously shown that maintenance of telomere length by ectopic hTERT expression does not prevent epigenetic ageing ^55^. Much remains to be explored in this intriguing area.

Transcription factors recognize and bind to unique and specific motifs on the genome to stimulate expression of down-stream genes. Methylation or demethylation of these motifs can alter the access, binding and eventual efficacy of the transcriptions factors in stimulating gene expression. Our analysis showed that CpGs within the TFAP2C-binding motif become increasingly methylated with age, which inhibits the binding of TFAP2C to its binding site ^56^. Increasing methylation of this motif was also identified in ageing cats, cattle and bats ^57–59^. Such inter-species concordance is precisely what is expected and sought after to help identify the common underlying mechanism of ageing across the mammalian class. How age-related decline in TFAP2C activity is linked to epigenetic ageing is presently unclear, but it might involve preservation of cell identity, as TFAP2C protein regulates the expression of genes which participate in development processes and interestingly, in concert with Gata3 and Eomes, TFAP2C induces a high-degree of nuclear reprogramming of fibroblasts into trophoblast stem cells ^56^. In a related way, the POU3F1 gene is also implicated by the proximal location of some age-related CpGs. The protein product of this gene, Oct6 is also able to elicit reprogramming of cells ^60^.

In keeping within the context of cellular identity, it is notable that CpGs proximal to ZNF536 becomes increasingly demethylated in ageing humans and elephants. This zinc-finger protein at higher levels, inhibit neuronal differentiation ^61^. The regulation of cellular identity is an important feature of development and it is consistent with gene ontology analysis which singled out genes related to the process of development and cellular differentiation to be highly implicated by age-related CpG methylation change. The involvement of differentiation and development have also been previously highlighted in similar analysis of human DNA methylation ^28^. Although detailed mechanistic understanding of how developmental processes are tied to epigenetic ageing remains unclear, their involvement is not in doubt, and in-depth investigation into this link is clearly warranted and holds great promise.

Our EWAS of elephant age implicated mTORC1 whose inhibition is known to increase the life span of yeast ^62^, worms ^63^, and mice ^64,65^. Recently, we demonstrated that inhibition of mTOR activity by rapamycin retarded epigenetic ageing of normal human cells ^66^. Collectively, these observations carry deep implications because the involvement of mTOR in the ageing of these different organisms (yeast to elephant) suggest that the conservation of ageing mechanism observed among mammals may extend to evolutionarily distant classes, and that the mechanism of ageing is a primitive one that emerged early on in life on earth.

While the above highlights underlying concordance of ageing among different species, there were also clear differences between age-related CpGs of elephants and humans. Methylation of CpGs proximal to TNF change in opposite directions between ageing humans and elephants. At first instance, this would appear to mean that TNF plays different roles with regards to ageing in these two species; which is possible but perhaps rather less likely. Predictions of downstream outcomes based on a single gene is fraught with uncertainties as almost all biological traits are caused by intricate network of proteins that mediate and regulate almost every step. Hence the same pathway can be activated or repressed at many different points along the entire network. An example that exhibits this apparent contradiction is RRAGC, which encodes a protein that is required for mTORC1 activity ^62^. The CpG proximal to RRAGC appears to be hypermethylated with age in elephants but unchanged in humans. At first sight, it would suggest that mTOR is not involved in ageing of human cells. Yet, we have demonstrated that inhibition of mTOR retards epigenetic ageing of human cells ^66^. Such apparent irregularity can be better understood by considering the fact that there are many ways to regulate activity of an enzyme or signaling pathway, and different species may employ different ways to do so. This highlights the challenge in attempting to comprehend the final consequence from the change of activity of a single gene. Therefore, while we can be fairly certain of the obvious similarities of age-related pathways between different species, the apparent divergent ones are much more difficult to interpret.

In summary, biological processes implicated in ageing and shared between elephants and humans include: cellular differentiation, organism development, metabolism and circadian rhythm. Shared age-related molecular changes are methylation of bivalent chromatin domain, targets of polycomb repressive complex and TFAP2C binding sites. Ultimately, it will be necessary to understand how these age-associated biological changes reduce fitness and health, which characterizes ageing of all animals.

### Experimental Procedures

#### Study Animals

Our study population included 140 elephants (57 African and 83 Asian) housed in 27 AZA-accredited zoos in North America (including Canada) (Table 1). Known or estimated birthdates were gleaned from each species’ studbooks. This study was authorized by the management of each participating zoo and, where applicable, was reviewed and approved by zoo research committees. In addition, the study received IACUC s#18-29) at the NZP; and endorsement from the elephant Taxon Advisory Group and Species Survival Plan.

#### Blood collection

Whole blood samples from either an ear or leg vein directly into an EDTA tube were collected between 1998 and 2019 during regular veterinary examinations and shipped frozen to the genetics lab at the Smithsonian Conservation Biology Institute Center for Conservation Genomics (SCBI-CCG). The samples were stored in an ultralow freezer (−80C) until DNA extraction.

#### DNA Extraction

DNA was extracted from 250 μl aliquots of whole blood with the BioSprint 96 DNA Blood Kit (Qiagen Corp.) using the BioSprint 96 robot in 96-well format. DNA yield was measured using a Qubit high sensitivity, double stranded DNA kit (Life Technologies), using 1μL of input DNA.

#### Human tissue samples

To build the human-elephant clock, we analyzed previously generated methylation data from 850 human tissue samples (adipose, blood, bone marrow, dermis, epidermis, heart, keratinocytes, fibroblasts, kidney, liver, lung, lymph node, muscle, pituitary, skin, spleen) from individuals whose ages ranged from 0 to 93 years. Tissue and organ samples were obtained from the National NeuroAIDS Tissue Consortium ^67^, blood samples from the Cape Town Adolescent Antiretroviral Cohort Study ^68^, and skin and other primary cells were provided by Kenneth Raj ^55^. Ethics approval (IRB#15-001454, IRB#16-000471, IRB#18-000315, IRB#16-002028).

### DNA methylation data

We generated DNA methylation data using the custom Illumina chip “HorvathMammalMethylChip40”. The mammalian methylation array provides high coverage (over thousand X) of highly conserved CpGs in mammals, but focuses only on 36k CpGs that are highly conserved across mammals. DNA methylation arrays were profiled using a custom Illumina methylation array (HorvathMammalMethylChip40) based on 38k CpG sites in highly conserved regions in mammals. Out of 38,000 probes, 2,000 were selected based on their utility for human biomarker studies: these CpGs, which were previously implemented in human Illumina Infinium arrays (EPIC, 450K) were selected due to their relevance for estimating age, blood cell counts, or the proportion of neurons in brain tissue. The remaining 35,988 probes were chosen to assess cytosine DNA methylation levels in mammalian species (Arneson, Ernst, Horvath, in preparation). The particular subset of species for each probe is provided in the chip manifest file can be found at Gene Expression Omnibus (GEO) at NCBI as platform GPL28271. The SeSaMe normalization method was used to define beta values for each probe ^69^.

#### Penalized Regression models

Details on the clocks (CpGs, genome coordinates) and R software code are provided in the Supplement. Penalized regression models were created with glmnet ^70^. We investigated models produced by both “elastic net” regression (alpha=0.5). The optimal penalty parameters in all cases were determined automatically by using a 10 fold internal cross-validation (cv.glmnet) on the training set. The alpha value for the elastic net regression was set to 0.5 (midpoint between Ridge and Lasso type regression) and was not optimized for model performance.

We performed a cross-validation scheme for arriving at unbiased (or at least less biased) estimates of the accuracy of the different DNAm based age estimators. One type consisted of leaving out a single sample (LOOCV) from the regression, predicting an age for that sample, and iterating over all samples. A critical step is the transformation of chronological age (the dependent variable). While no transformation was used for the blood clock for elephants, we did use a log linear transformation of age for the dual species clock of chronological age.

#### Relative age estimation

To introduce biological meaning into age estimates of elephants and humans that have very different lifespans, as well as to overcome the inevitable skewing due to unequal distribution of data points from elephants and humans across age ranges, relative age estimation was made using the formula: Relative age= Age/maxLifespan, where the maximum lifespan for the two species was chosen from the anAge data base ^71^.

#### Epigenome wide association studies of age

EWAS was performed in each tissue separately using the R function “standardScreeningNumericTrait” from the “WGCNA” R package ^72^. Next the results were combined across tissues using Stouffer’s meta-analysis method. The analysis was done using the genomic region of enrichment annotation tool ^73^. The gene level enrichment was done using GREAT analysis and human Hg19 background ^73^.

#### Gene ontology enrichment analysis

The analysis was done using the genomic region of enrichment annotation tool ^47^. The gene level enrichment was done using GREAT analysis ^47^ and human Hg19 background. The background probes were limited to 21,377 probes that were mapped to the same gene in the African elephant genome. The top three enriched datasets from each category (Canonical pathways, diseases, gene ontology, human and mouse phenotypes, and upstream regulators) were selected and further filtered for significance at p < 10^−4^.

## Acknowledgements

This work was supported by the Paul G. Allen Frontiers Group (SH). We acknowledge SCBI intern Bethany Nelson, Karen Steinman (SPL) and the people from the following zoos for support of the project: Albuquerque Biological Park, Birmingham Zoo, Busch Gardens Tampa, Buttonwood Park Zoo, Calgary Zoo, Columbus Zoo, Dallas Zoo, D Animal Kingdom, Fresno Chaffee Zoo, Grants Farm, Houston Zoological Gardens, Indianapolis Zoological Society, Inc., Los Angeles Zoo and Botanical Gardens, National Zoological Park, Oregon Zoo, Roger Williams Zoo, Rosamond Gifford Zoo, San Diego Zoo, Seneca Park Zoo, Tennessee Elephant Sanctuary, Topeka Zoo, Toledo Zoo, Tulsa Zoo, Utah’s Hogle Zoo, and Zoo Miami.

## Funding

This work was supported by the Paul G. Allen Frontiers Group (SH). Stipend and reagent support for NAP was provided through Dr. Janice Sanders and the George E. Burch Post-Doctoral Fellowship in Theoretical Medicine and Affiliated Theoretical Science. The funders had no role in the study design, data collection, decision to publish, or preparation of the manuscript.

## Conflict of Interest Statement

SH is a founder of the non-profit Epigenetic Clock Development Foundation which plans to license several patents from his employer UC Regents. These patents list SH as inventor. The other authors declare no conflicts of interest.

## Author Contributions

NAP: DNA sample processing (DNA extraction and quantification). Contribution of data, DNA samples and data base management: NAP, JLB, SH, LB, JEM, MGC. Statistical analysis: JAZ, AH, MY, JE, SH. Drafting and editing the article: SH, NAP, JLB, KR. All authors participated in writing the article. SH conceived of the study.

## References

1 Lee, P. C., Sayialel, S., Lindsay, W. K. & Moss, C. J. African elephant age determination from teeth: validation from known individuals. African Journal of Ecology 50, 9–20, doi:10.1111/j.1365-2028.2011.01286.x (2012).

2 Lahdenperä, M., Mar, K. U. & Lummaa, V. Reproductive cessation and post-reproductive lifespan in Asian elephants and pre-industrial humans. Front Zool 11, 54, doi:10.1186/s12983-014-0054-0 (2014).

3 Hakeem, A. Y. et al. Brain of the African elephant (Loxodonta africana): neuroanatomy from magnetic resonance images. Anat Rec A Discov Mol Cell Evol Biol 287, 1117–1127, doi:10.1002/ar.a.20255 (2005).

4 Brown, J. L. in Reproductive Sciences in Animal Conservation 243–273 (Springer, 2019).

5 Byrne, R. W., Bates, L. A. & Moss, C. J. Elephant cognition in primate perspective. Comparative Cognition & Behavior Reviews 4, 65–79, doi:10.3819/ccbr.2009.40009 (2009).

6 Lee, P. C., Fishlock, V., Webber, C. E. & Moss, C. J. The reproductive advantages of a long life: longevity and senescence in wild female African elephants. Behav Ecol Sociobiol 70, 337–345, doi:10.1007/s00265-015-2051-5 (2016).

7 de Silva, S. et al. Demographic variables for wild Asian elephants using longitudinal observations. PLoS One 8, e82788, doi:10.1371/journal.pone.0082788 (2013).

8 Abegglen, L. M. et al. Potential Mechanisms for Cancer Resistance in Elephants and Comparative Cellular Response to DNA Damage in Humans. JAMA 314, 1850–1860, doi:10.1001/jama.2015.13134 (2015).

9 Sukumar, R., Joshi, N. & Krishnamurthy, V. Growth in the Asian elephant. Proceedings: Animal Sciences 97, 561–571 (1988).

10 Arivazhagan, C. & Sukumar, R. Constructing Age Structures of Asian Elephant Populations: A Comparison of Two Field Methods of Age Estimation. Gajah 29(2008).

11 Laws, R. Age criteria for the African elephant: Loxodonta a. africana. African Journal of Ecology 4, 1–37 (1966).

12 Jachmann, H. Estimating age in African elephants: a revision of Laws’ molar evaluation technique. African Journal of Ecology 26, 51–56, doi:10.1111/j.1365-2028.1988.tb01127.x (1988).

13 Roth, V. L. & Shoshani, J. Dental identification and age determination in Elephas maximus. Journal of Zoology 214, 567–588, doi:10.1111/j.1469-7998.1988.tb03760.x (1988).

14 Lang, E. M. Observations on growth and molar change in the African elephant. African Journal of Ecology 18, 217–234, doi:10.1111/j.1365-2028.1980.tb00643.x (1980).

15 Sikes, S. K. The African elephant, Loxodonta africana: a field method for the estimation of age. Journal of Zoology 154, 235–248, doi:10.1111/j.1469-7998.1968.tb01661.x (1968).

16 Laws, R. M. Eye Lens Weight And Age In African Elephants. African Journal of Ecology 5, 46–52, doi:10.1111/j.1365-2028.1967.tb00760.x (1967).

17 Laws, R. M., Parker, I. S. C. & Archer, A. L. Estmating Ltve Weights Of Eijephants From Hind Leg Weights. African Journal of Ecology 5, 106–111, doi:10.1111/j.1365-2028.1967.tb00765.x (1967).

18 Pilgram, T. & Western, D. Inferring the sex and age of African elephants from tusk measurements. Biological Conservation 36, 39–52, doi:https://doi.org/10.1016/0006-3207(86)90100-X (1986).

19 Shrader, A. M. et al. Growth and age determination of African savanna elephants. Journal of Zoology 270, 40–48, doi:10.1111/j.1469-7998.2006.00108.x (2006).

20 Kongrit, C. & Siripunkaw, C. Determination of age and construction of population age structure of wild Asian elephants based on dung bolus circumference. The Thai Journal of Veterinary Medicine 47, 145–153 (2017).

21 Lark, R. M. A comparison between techniques for estimating the ages of African elephants (Loxodonta africana). African Journal of Ecology 22, 69–71, doi:10.1111/j.1365-2028.1984.tb00678.x (1984).

22 Stansfield, F. J. A Novel Objective Method of Estimating the Age of Mandibles from African Elephants (Loxodonta africana Africana). PLOS ONE 10, e0124980, doi:10.1371/journal.pone.0124980 (2015).

23 Smith, Z. D. & Meissner, A. DNA methylation: roles in mammalian development. Nat Rev Genet 14, 204–220 (2013).

24 Rakyan, V. K. et al. Human aging-associated DNA hypermethylation occurs preferentially at bivalent chromatin domains. Genome research 20, 434–439, doi:10.1101/gr.103101.109 (2010).

25 Teschendorff, A. E. et al. Age-dependent DNA methylation of genes that are suppressed in stem cells is a hallmark of cancer. Genome research 20, 440–446, doi:10.1101/gr.103606.109 (2010).

26 Horvath, S. & Raj, K. DNA methylation-based biomarkers and the epigenetic clock theory of ageing. Nat Rev Genet, doi:10.1038/s41576-018-0004-3 (2018).

27 Field, A. E. et al. DNA Methylation Clocks in Aging: Categories, Causes, and Consequences. Mol Cell 71, 882–895, doi:10.1016/j.molcel.2018.08.008 (2018).

28 Horvath, S. DNA methylation age of human tissues and cell types. Genome Biol 14, R115, doi:10.1186/gb-2013-14-10-r115 (2013).

29 Marioni, R. et al. DNA methylation age of blood predicts all-cause mortality in later life. Genome Biol. 16, 25 (2015).

30 Christiansen, L. et al. DNA methylation age is associated with mortality in a longitudinal Danish twin study. Aging Cell 15, 149–154, doi:10.1111/acel.12421 (2016).

31 Perna, L. et al. Epigenetic age acceleration predicts cancer, cardiovascular, and all-cause mortality in a German case cohort. Clin Epigenetics 8, 64, doi:10.1186/s13148-016-0228-z (2016).

32 Chen, B. H. et al. DNA methylation-based measures of biological age: meta-analysis predicting time to death. Aging (Albany NY) 8, 1844–1865, doi:10.18632/aging.101020 (2016).

33 Horvath, S. & Levine, A. J. HIV-1 Infection Accelerates Age According to the Epigenetic Clock. J Infect Dis 212, 1563–1573, doi:10.1093/infdis/jiv277 (2015).

34 Marioni, R. E. et al. The epigenetic clock is correlated with physical and cognitive fitness in the Lothian Birth Cohort 1936. Int J Epidemiol 44, 1388–1396, doi:10.1093/ije/dyu277 (2015).

35 Levine, M. E., Lu, A. T., Bennett, D. A. & Horvath, S. Epigenetic age of the pre-frontal cortex is associated with neuritic plaques, amyloid load, and Alzheimer’s disease related cognitive functioning. Aging (Albany NY) 7, 1198–1211, doi:10.18632/aging.100864 (2015).

36 Horvath, S. et al. Accelerated epigenetic aging in Down syndrome. Aging Cell 14, 491–495, doi:10.1111/acel.12325 (2015).

37 Maierhofer, A. et al. Accelerated epigenetic aging in Werner syndrome. Aging (Albany NY) 9, 1143–1152, doi:10.18632/aging.101217 (2017).

38 Horvath, S. et al. Huntington’s disease accelerates epigenetic aging of human brain and disrupts DNA methylation levels. Aging (Albany NY) 8, 1485–1512, doi:10.18632/aging.101005 (2016).

39 Horvath, S. et al. Obesity accelerates epigenetic aging of human liver. Proc Natl Acad Sci U S A 111, 15538–15543, doi:10.1073/pnas.1412759111 (2014).

40 Levine, M. E. et al. Menopause accelerates biological aging. Proc Natl Acad Sci U S A 113, 9327–9332, doi:10.1073/pnas.1604558113 (2016).

41 Fahy, G. M. et al. Reversal of epigenetic aging and immunosenescent trends in humans. Aging Cell 18, e13028, doi:10.1111/acel.13028 (2019).

42 Bell, C. G. et al. DNA methylation aging clocks: challenges and recommendations. Genome Biology 20, 249, doi:10.1186/s13059-019-1824-y (2019).

43 Petkovich, D. A. et al. Using DNA Methylation Profiling to Evaluate Biological Age and Longevity Interventions. Cell Metab 25, 954–960 e956, doi:10.1016/j.cmet.2017.03.016 (2017).

44 Cole, J. J. et al. Diverse interventions that extend mouse lifespan suppress shared age-associated epigenetic changes at critical gene regulatory regions. Genome Biol 18, 58, doi:10.1186/s13059-017-1185-3 (2017).

45 Wang, T. et al. Epigenetic aging signatures in mice livers are slowed by dwarfism, calorie restriction and rapamycin treatment. Genome Biol 18, 57, doi:10.1186/s13059-017-1186-2 (2017).

46 Stubbs, T. M. et al. Multi-tissue DNA methylation age predictor in mouse. Genome Biol 18, 68, doi:10.1186/s13059-017-1203-5 (2017).

47 Thompson, M. J. et al. A multi-tissue full lifespan epigenetic clock for mice. Aging (Albany NY) 10, 2832–2854, doi:10.18632/aging.101590 (2018).

48 Meer, M. V., Podolskiy, D. I., Tyshkovskiy, A. & Gladyshev, V. N. A whole lifespan mouse multi-tissue DNA methylation clock. eLife 7, e40675, doi:10.7554/eLife.40675 (2018).

49 Bruunsgaard, H., Skinhoj, P., Pedersen, A. N., Schroll, M. & Pedersen, B. K. Ageing, tumour necrosis factor-alpha (TNF-alpha) and atherosclerosis. Clin Exp Immunol 121, 255–260, doi:10.1046/j.1365-2249.2000.01281.x (2000).

50 Bauernfeind, F., Niepmann, S., Knolle, P. A. & Hornung, V. Aging-Associated TNF Production Primes Inflammasome Activation and NLRP3-Related Metabolic Disturbances. J Immunol 197, 2900–2908, doi:10.4049/jimmunol.1501336 (2016).

51 de Flamingh, A., Coutu, A., Roca, A. L. & Malhi, R. S. Accurate Sex Identification of Ancient Elephant and Other Animal Remains Using Low-Coverage DNA Shotgun Sequencing Data. G3 (Bethesda) 10, 1427–1432, doi:10.1534/g3.119.400833 (2020).

52 Wilcox, A. G., Vizor, L., Parsons, M. J., Banks, G. & Nolan, P. M. Inducible Knockout of Mouse Zfhx3 Emphasizes Its Key Role in Setting the Pace and Amplitude of the Adult Circadian Clock. J Biol Rhythms 32, 433–443, doi:10.1177/0748730417722631 (2017).

53 White, A. J., Kresovich, J. K., Xu, Z., Sandler, D. P. & Taylor, J. A. Shift work, DNA methylation and epigenetic age. Int J Epidemiol 48, 1536–1544, doi:10.1093/ije/dyz027 (2019).

54 Lu, A. T. et al. GWAS of epigenetic aging rates in blood reveals a critical role for TERT. Nat Commun 9, 387, doi:10.1038/s41467-017-02697-5 (2018).

55 Kabacik, S., Horvath, S., Cohen, H. & Raj, K. Epigenetic ageing is distinct from senescence-mediated ageing and is not prevented by telomerase expression. Aging (Albany NY) 10, 2800–2815, doi:10.18632/aging.101588 (2018).

56 Woodfield, G. W., Hitchler, M. J., Chen, Y., Domann, F. E. & Weigel, R. J. Interaction of TFAP2C with the estrogen receptor-alpha promoter is controlled by chromatin structure. Clinical cancer research: an official journal of the American Association for Cancer Research 15, 3672–3679, doi:10.1158/1078-0432.ccr-08-2343 (2009).

57 Raj, K. et al. Epigenetic clock and methylation studies in cats. bioRxiv, 2020.2009.2006.284877, doi:10.1101/2020.09.06.284877 (2020).

58 Kordowitzki, P. et al. Epigenetic clock and methylation study of oocytes from a bovine model of reproductive aging. bioRxiv, 2020.2009.2010.290056, doi:10.1101/2020.09.10.290056 (2020).

59 Wilkinson, G. S. et al. Genome Methylation Predicts Age and Longevity of Bats. bioRxiv, 2020.2009.2004.283655, doi:10.1101/2020.09.04.283655 (2020).

60 Kim, K. P. et al. Reprogramming competence of OCT factors is determined by transactivation domains. Sci Adv 6, doi:10.1126/sciadv.aaz7364 (2020).

61 Qin, Z. et al. ZNF536, a novel zinc finger protein specifically expressed in the brain, negatively regulates neuron differentiation by repressing retinoic acid-induced gene transcription. Mol Cell Biol 29, 3633–3643, doi:10.1128/mcb.00362-09 (2009).

62 Yang, G. et al. RagC phosphorylation autoregulates mTOR complex 1. EMBO J 38, doi:10.15252/embj.201899548 (2019).

63 Robida-Stubbs, S. et al. TOR signaling and rapamycin influence longevity by regulating SKN-1/Nrf and DAF-16/FoxO. Cell Metab 15, 713–724, doi:10.1016/j.cmet.2012.04.007 (2012).

64 Zhang, Y. et al. Rapamycin extends life and health in C57BL/6 mice. J Gerontol A Biol Sci Med Sci 69, 119–130, doi:10.1093/gerona/glt056 (2014).

65 Weichhart, T. mTOR as Regulator of Lifespan, Aging, and Cellular Senescence: A Mini-Review. Gerontology 64, 127–134, doi:10.1159/000484629 (2018).

66 Horvath, S., Lu, A. T., Cohen, H. & Raj, K. Rapamycin retards epigenetic ageing of keratinocytes independently of its effects on replicative senescence, proliferation and differentiation. Aging (Albany NY) 11, 3238–3249, doi:10.18632/aging.101976 (2019).

67 Morgello, S. et al. The National NeuroAIDS Tissue Consortium: a new paradigm in brain banking with an emphasis on infectious disease. Neuropathol Appl Neurobiol 27, 326–335. (2001).

68 Horvath, S. et al. Perinatally acquired HIV infection accelerates epigenetic aging in South African adolescents. AIDS (London, England) 32, 1465–1474, doi:10.1097/QAD.0000000000001854 (2018).

69 Zhou, W., Triche, T. J., Jr, Laird, P. W. & Shen, H. SeSAMe: reducing artifactual detection of DNA methylation by Infinium BeadChips in genomic deletions. Nucleic Acids Research 46, e123–e123, doi:10.1093/nar/gky691 (2018).

70 Friedman, J., Hastie, T. & Tibshirani, R. Regularization Paths for Generalized Linear Models via Coordinate Descent. Journal of Statistical Software 33, 1–22 (2010).

71 de Magalhaes, J. P., Costa, J. & Church, G. M. An analysis of the relationship between metabolism, developmental schedules, and longevity using phylogenetic independent contrasts. J Gerontol A Biol Sci Med Sci 62, 149–160 (2007).

72 Langfelder, P. & Horvath, S. WGCNA: an R package for weighted correlation network analysis. BMC Bioinformatics 9, 559 (2008).

73 McLean, C. Y. et al. GREAT improves functional interpretation of cis-regulatory regions. Nat Biotechnol 28, doi:10.1038/nbt.1630 (2010).

74 McLean, C. Y. et al. GREAT improves functional interpretation of cis-regulatory regions. Nat Biotechnol 28, 495–501, doi:10.1038/nbt.1630 (2010).

